# Principles of 3D Nucleus Organization and Epigenetic Regulation in Diploid Genome Revealed by Multi-omic Data from Hybrid Mouse

**DOI:** 10.1101/783662

**Authors:** Zhijun Han, Cui Kairong, Katarzyna Placek, Ni Hong, Chengqi Lin, Wei Chen, Keji Zhao, Wenfei Jin

**Affiliations:** Department of Biology, Southern University of Science and Technology, Shenzhen, Guangdong 518055, China; Institute of Life Sciences, Southeast University, Nanjing 210096, China; Systems Biology Center, National Heart, Lung and Blood Institute, NIH, Bethesda, MD 20892, USA

**Keywords:** Hybrid mouse, 3D nucleus, chromatin architecture, Hi-C, epigenetic regulations

## Abstract

Most mammalian genomes are diploid and previous studies have extensively investigated the average epigenetic profiles of homologous chromosomes. Here we use hybrid mice to distinguish the epigenetic status and three-dimensional organization of homologous chromosomes. We generated Hi-C, ChIP-seq and RNA-seq datasets from CD4 T cells of B6, Cast and hybrid mice, respectively, and systematically analyzed the 3D nucleus organization and epigenetic regulation. Our data indicate that the inter-chromosomal interaction patterns between homologous chromosomes are similar and the similarity is highly correlated with their allelic co-expression levels. Construction of 3D nucleus based on allele-specific interaction frequency revealed symmetric positioning of homologous chromosomes in the 3D nuclear space. The inter-chromosomal interactions at centromeres are significantly weaker than those at telomeres, indicating positioning of centromeres toward the inside of chromosome territories and telomeres toward the surface of chromosome territories. The majority A|B compartments or topologically associated domains (TADs) are consistent between B6 and Cast. We found 58% of the haploids in hybrids maintain their parental compartment status at B6/Cast divergent compartments due to cis-effect. About 95% of the trans-effected B6/Cast divergent compartments converge to same compartment status potentially due to a shared cellular environment. We found the differentially expressed genes between the two haploids in hybrid were associated with either genetic associated cis-effects or epigenetic associated trans-effects. The widespread epigenetic differences between B6 and Cast suggest that epigenetic changes may be major contributors to differences between B6 and Cast. Our data revealed symmetrical positioning of homologous chromosomes in 3D nucleus and enhanced our understanding of allele-specific epigenetic regulation.

## Introduction

Chromatin is well organized within 3D nucleus, which renders efficient packaging of DNA while simultaneously allowing precise gene regulation and genome replication. Although study of DNA folding and packaging in nucleus has fascinated biologists for over a century, the higher-order chromatin organization in nucleus and its influence on gene expression is still not fully understood. Studies based on Hi-C showed that the chromatin is segregated into A and B compartments, which correspond to euchromatin and heterochromatin, respectively (Lieberman-Aiden et al. 2009). While A compartments are more accessible to DNase I, more gene rich, more transcriptionally active, and is associated with higher active histone enrichment, B compartments are generally associated with transcriptional repression. Furthermore, mammalian genome is organized into a large number of topologically associated domains (TADs) (Dixon et al. 2012) and active TADs appear to be segregated from inactive TADs (Simonis et al. 2006; Lieberman-Aiden et al. 2009; de Wit et al. 2013; Gibcus and Dekker 2013; Meuleman et al. 2013; Naumova et al. 2013), which is conserved in *Drosophila* (Sexton et al. 2012). The TAD structure is relatively stable across different cell types, which requires the function of CTCF and cohesin (Dixon et al. 2012; Zuin et al. 2014; Nora et al. 2017; Ren et al. 2017). It has been shown that disruption of TAD structure leads to various diseases (Franke et al. 2016; Lupianez et al. 2016).

Most mammalian genomes are diploid, one is paternal and the other is maternal. Previous studies have provided rich information about the dynamics of chromatin status and histone modification during the early gamete and embryo development (Liu et al. 2016; Lu et al. 2016; Jung et al. 2017; Nagano et al. 2017; Stevens et al. 2017). However, almost all the studies only inferred the average chromatin interactions and average epigenetic status across the diploid genomes, partially because genetic variations in humans or mice could not provide sufficient segregating sites for investigating the allele-specific interaction and haploid-specific chromatin status. Although recent studies by Tan et al. investigated the diploid genome at single cell resolution (Tan et al. 2018), characterizing the 3D genome structures of diploid mammalian cells remains challenging and the principle of 3D nucleus organization is a mystery to some extent (Carstens et al. 2016).

In this study, we introduced a hybrid mouse system with paternal B57CL/6J (B6) and maternal CAST/EiJ (Cast) that contained a large number of segregating sites for distinguishing paternal haploid from maternal haploid, which serves as an ideal model for investigating the allele-specific 3D nucleus organization and epigenetic regulation. We used naïve CD4 T cells from the hybrid mice and their parents to generate multi-omics datasets including 3eHi-C, ChIP-seq and RNA-seq. Interestingly, we found the inter-chromosomal interaction pattern of each chromosome was highly similar to that of its homologous counterpart. In particular, the similarity of interaction pattern is highly correlated with allelic co-expression level. We further found that positioning of homologous chromosomes in 3D nucleus is symmetrical; centromeres are likely to be located near the center of chromosome territories while telomeres tend to be at the surface of chromosome territories. Our data indicates that 42% haploids at B6/Cast divergent compartments converge to the same A|B compartments in hybrid due to shared cellular environment or trans-effect. We found that haploid-specific TAD boundaries and histone modifications are correlated with haploid specific expression. Epigenetic differences between B6 and Cast are much higher than that of genetic differences, indicating epigenetic changes are major contributors of B6 and Cast differences.

## Results

### Constructing the hybrid mouse system and generating multi-omics datasets

Hybrid carries the homologous chromosomes from two different populations or species and its genome carries millions of segregating single nucleotide polymorphisms (SNPs). Therefore, hybrid provides an ideal model for investigating bi-parentally inheritable traits and allele-specific regulation of gene expression. For gaining insights into the distinct 3D nucleus organization and epigenetic status between the two homologous chromosomes, we constructed a hybrid mouse system by crossbreeding male C57BL/6J (B6) and female CAST/EiJ (Cast) mice (Fig 1A). Naïve CD4 T cells from 6-8 weeks mice were isolated for systematic analysis of 3D nucleus organization, epigenetic status, and gene expression of the two distinct parental alleles in the hybrid mice. We applied the 3e Hi-C protocol (Ren et al. 2017) to capture the global chromatin interaction information, ChIP-seq to profile genome-wide histone modification (or transcription factor binding), and RNA-seq to profile gene expression in both parental mice and hybrid mice.

**Fig 1.**
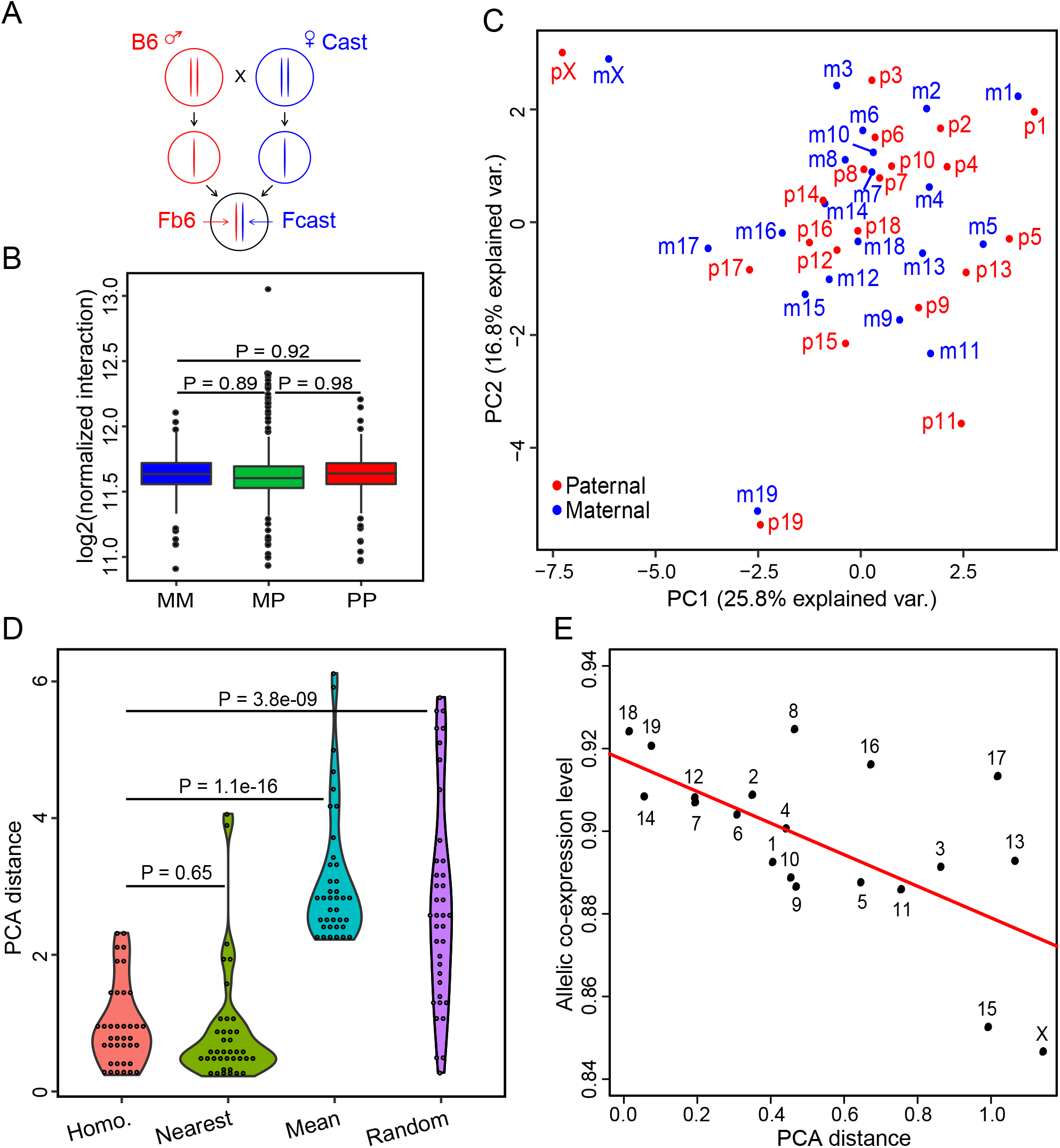
Homologous chromosomes showed similar interaction patterns and the similarity correlate with allelic co-expressions level. Abbreviation: p, paternal chromosomes; m, maternal chromosomes. **A)** The hybrid mouse model system. Male C57BL/6J (B6) and female CAST/EiJ (Cast) mice were crossbred to generate hybrid mouse. In the hybrid mouse, haploid genome originating from B6 and Cast were called Fb6 and Fcast, respectively. **B)** Boxplot of interaction intensities between maternal-maternal (MM), maternal-paternal (MP) and paternal-paternal (PP) chromosome pairs based on allelic specific PETs. The interactions intensities among MM, MP and PP did not show any significant differences. P values were calculated using t test. **C)** PCA analysis of chromosomal interactions. The chromosomes are projected on 2D plot, in which short distance indicated high similarity of interaction patterns. **D)** Distance between homologous chromosomes almost always are the shortest among all potential pairs on the 2D PCA projection. Abbreviations: Homo, distance between homologous chromosome pair; nearest, distance between a chromosome and its nearest non-homologous chromosome; mean, mean distance between a chromosome and all the other chromosomes; random, distance between two randomly picked chromosomes. A short distance indicates a high similarity of interaction pattern. P values were calculated using t test. **E)** Bi-allelic co-expression level is correlated with the similarity of chromosomal interaction pattern between homologous chromosomes. (R = 0.7, P value = 0.0006)

In order to investigate the chromatin interaction at haploid level, we assigned each end of the paired-end-tags (PETs) to the B6 haploid and Cast haploid, respectively, based on the strain-specific allele on the PETs (see Methods). For convenience, haploids in hybrid mice inherited from B6 and Cast were called Fb6 and Fcast, respectively. The length distribution of PETs in Fb6 and Fcast are similar to that in B6 or Cast Hi-C libraries (Fig S1A).

### Homologous chromosomes exhibit similar interaction patterns

The millions of segregating sites in hybrid mice allowed us to distinguish the two haploids and to construct diploid 3D nucleus. We built an inter-chromosomal interaction matrix at haploid level using the Hi-C PETs with both ends having strain-specific alleles. We classified the inter-chromosomal interactions into three groups, namely maternal-maternal (MM), maternal-paternal (MP) and paternal-paternal (PP). Our analysis showed the normalized interaction densities among MM, MP and PP did not have any significant difference (Fig 1B), indicating the chromosomes’ paternal or maternal origination do not bias genome-wide interaction pattern.

We conducted principal component analysis (PCA) on the haploid-level inter-chromosomal interaction matrix to capture the principles underlying inter-chromosomal interactions. We found the homologous chromosomes are in close proximity on the PCA projection compared to non-homologous chromosomes (Fig 1C), potentially indicating the homologous chromosomes have similar interaction patterns. We further calculated the distance between any two chromosomes on PCA projection, namely PCA distance, to indicate the similarity of interaction patterns. The PCA distance between homologous chromosomes is comparable to distance between two nearest non-homologous chromosomes, and significantly shorter than the mean distance of a chromosome to other chromosomes, as well as significantly shorter than the distance between two random chromosomes (Fig 1D), further indicating that the similarity of interaction patterns between homologous chromosomes is one of the major features in genome-wide inter-chromosomal interactions. We further masked the interactions between homologous chromosomes and conducted PCA. The homologous chromosomes are still near each other on the PCA projection (Fig S2A), indicating the similarity of interaction patterns between homologous chromosomes is not caused by PETs directly linked them.

It is well known that mammalian female has two copies of X chromosome, with one X chromosome randomly silenced due to the dosage compensation effects (Schulz and Heard 2013; Crane et al. 2015). Both paternal and maternal X chromosomes are far from the autosomes in the PCA projection (Fig 1C), indicating the interaction patterns of X chromosome are quite different from that of autosomes. The result is consistent with a recent report that the imprinted X chromosome was highly compacted and had very few interactions with other chromosomes (Wang et al. 2016). Recent studies have shown that the imprinted X chromosome was separated into active particle (X-a) and inactive particle (X-i) by Dxz4 locus (Deng et al. 2015; Marks et al. 2015). We found the similarity of interaction pattern between homologous X-a was higher than that of X-i (Fig S2B), which is consistent with our expectation.

### Similarity of interaction patterns between homologous chromosomes is correlated with bi-allelic co-expression level

The overall expression level of a gene needs coordinated regulation between the two homologous alleles in diploid mammalian cells. However, whether the interaction pattern of homologous chromosomes affects the co-expression of bi-allelic genes is not clear. We calculated the bi-allelic co-expression level of each homologous chromosome pair by calculating correlation coefficient of allele-specific expressions between homologous chromosomes. We found that the similarity of interaction patterns between homologous chromosomes is highly correlated with the bi-allelic co-expression level (Fig 1E, R = 0.7, P value = 0.0006), potentially indicating that chromosomal interaction may affect the bi-allelic co-expression. Since X-a and X-i exhibit different characters and properties (Deng et al. 2015; Marks et al. 2015), we separated X chromosome into X-a and X-i and analyzed their differences. In the PCA projection, the X-a exhibits autosome-like pattern, with both high similarity of interaction pattern and high allelic co-expression level, while the X-i showed low interaction similarity and low allelic co-expressions and thus far separated from autosomes (Fig S2C, R = 0.66, P value = 0.001).

### Spatial organization of haploid chromosomes in the nucleus

Chromosome territories have been studied by various chromosome imaging techniques (Cremer et al. 2003; Bolzer et al. 2005; Hua and Mikawa 2018). Interestingly, these studies found that homologous chromosomes are separated from each other in the nuclear space with strong cellular variation. Yet it is difficult to find a general trend of relative positioning of the homologous chromosomes in the nucleus due to a limited number of cells in these studies. In contrast, Hi-C data are collected from millions of cells and thus provide information on the average positioning of homologous chromosomes in the nucleus. We found homologous chromosomes have highly similar interaction patterns compared to non-homologous chromosomes, and the similarity of interaction patterns is highly correlated with the bi-allelic co-expression of the homologous chromosomes. To resolve the relative positioning of homologous chromosomes in nucleus, we constructed 3D nucleus based on the allele-specific inter-chromosomal interactions matrix. We built an iteratively weighted adjusting model to fit each haploid chromosome into the 3D nuclear space from the allele-specific interaction matrix (Fig S3A; see Methods). In our constructed 3D nucleus, the large chromosomes tended to locate in one polar and the small chromosomes locate in the other polar (Fig 2A). The paternal and maternal X chromosomes are far away from each other, consistent with a recent study (Giorgetti et al. 2016). We also found that chromosomes 19 are located on the surface of nucleus (Fig 2A), which is consistent with the constructed 3D nucleus in mouse embryonic stem (ES) cells without allelic information (Stevens et al. 2017).

**Figure 2.**
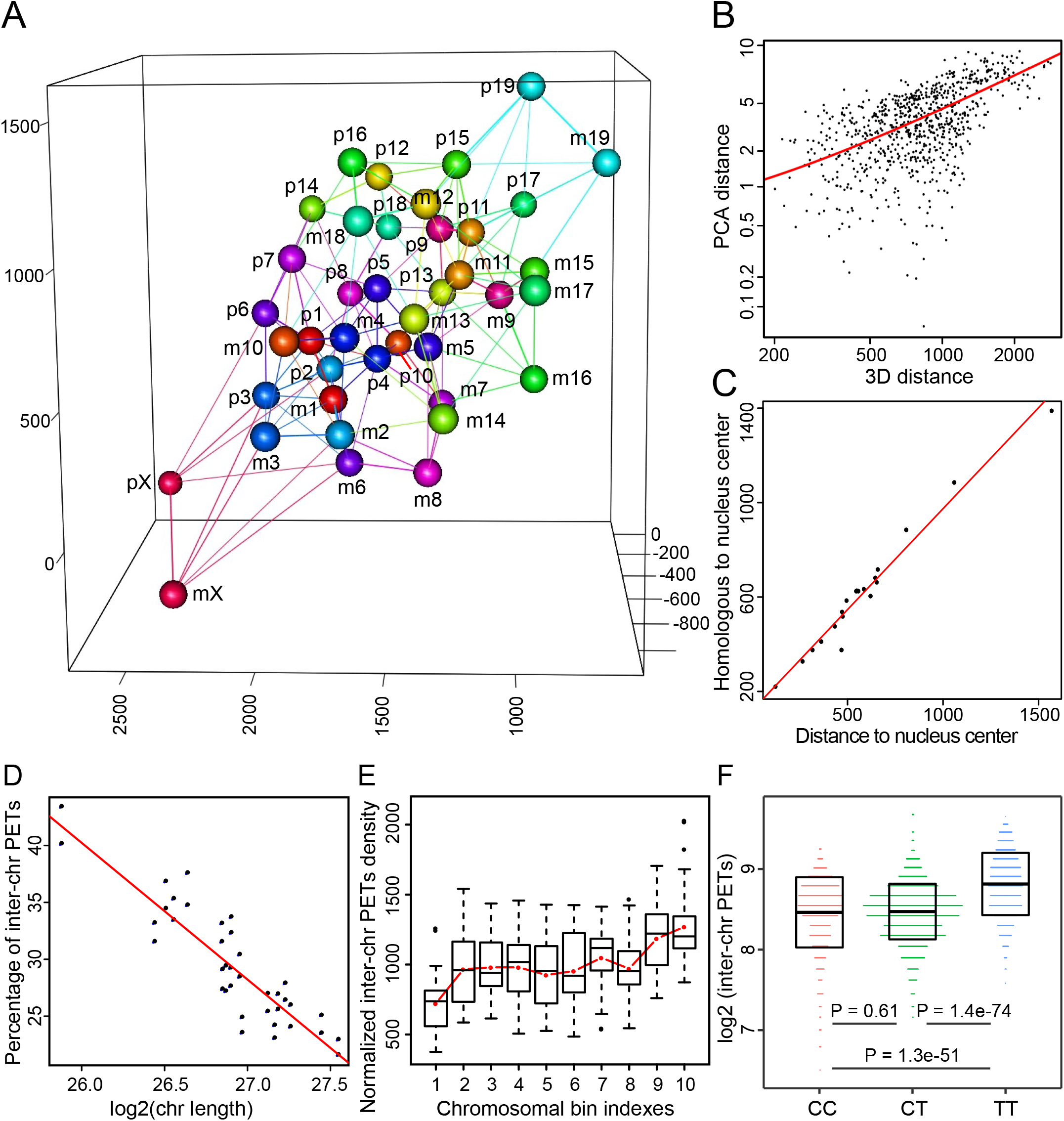
The 3D nucleus and principles of nucleus organization. A) Constructed 3D nucleus at chromosomal resolution. Each chromosome and it’s five closest chromosomes are connected by lines. Abbreviation: p, paternal chromosomes; m, maternal chromosomes. B) Chromosomal interaction pattern is positively correlated with chromosomal distance in the 3D nucleus. (R = 0.67, P value < 2.2e-16) C) Distances to center of 3D nucleus for paternal chromosomes are highly correlated with that of their maternal homologous counterpart. Center of 3D nucleus is calculated as the mean coordinate of all chromosomes. (R = 0.98, P value = 1.2e-14) D) The lengths of chromosomes are negatively correlated with the percentages of inter-chromosomal PETs. (R = 0.9, P value = 1.5e-14) E) Fractions of inter-chromosomal PETs are increasing from centromere to telomere along the chromosomes. Each chromosome is evenly divided into 10 bins. Red line plots the mean value for each bin. F) Telomere ends have much stronger inter-chromosomal interactions than that of centromere ends. Each chromosome is evenly binned into three parts, namely centromere end (C), middle (ignored) and telomere end (T). P values were calculated using unpaired t test.

We further found the inter-chromosomal distance in 3D nucleus correlated with PCA distance that represents interaction patterns (Fig 2B, R = 0.67, P value < 2.2e-16), which is consistent with the notion that the positioning of a chromosome in 3D nucleus contributes critically to chromatin interaction patterns. Interestingly, we noticed that homologous chromosomes seem to be positioned symmetrically surrounding the center of the 3D nucleus. We inferred the center of the 3D nucleus according to the coordinates of all chromosomes. Interestingly, we found the distance of a chromosome to the center of 3D nucleus is highly correlated with that of its homologous counterpart (Fig 2C, R = 0.98, P value = 1.2e-14), which strongly supports the notion that homologous chromosomes are symmetrically positioned surrounding the center of the 3D nuclear space. We also discovered the distance of a chromosome to paternal and maternal homologous chromosome is highly correlated (Fig S3B, R = 0.87, P value < 2.2e-16). In fact, the symmetric distribution of homologous chromosomes in nucleus could explain many biological phenomena including the similarity of inter-chromosomal interaction patterns.

It is well known that each chromosome occupies a spatially limited, roughly elliptical region termed as chromosome territory (Cremer and Cremer 2001; Meaburn and Misteli 2007). The lengths of chromosome are negatively correlated with the percentages of inter-chromosomal PETs (Fig 2D, R = 0.9, P value = 1.5e-14), which implies larger chromosomes have lower percentage of inter-chromosomal interactions than smaller chromosomes. These results indicate that the longer the chromosome, the bigger the chromosome territory, thus the smaller surface-area-to-volume ratio and lower fraction of inter chromosomes interactions. To determine the relative position of centromere and telomere within the chromosome territory, we evenly separated each chromosome from centromere to telomere into 10 bins and fraction of inter-chromosomal PETs in each bin is calculated. We found the fractions of inter-chromosomal PETs are increasing from centromere to telomere along the chromosome (Fig 2E), indicating that the centromere-end is more likely located in the center of a 3D chromosome territory while the telomere-end is more likely located at the surface of the 3D chromosome territory. We further calculated the interactions of centromere-centromere (CC), centromere-telomere (CT) and telomere-telomere (TT). Indeed, we found TT interactions are significantly higher than of CT and CC, although CC interactions and CT interactions are not significantly different (Fig 2F), which strongly supports our hypothesis that centromere are located in the center of each 3D chromosome territory.

### Parental divergent compartments transit into same status in hybrid mice

A|B compartments are generally in mega-base scale and functionally separate chromosomal regions (Lieberman-Aiden et al. 2009; Fortin and Hansen 2015). Global A|B compartment score analysis showed the chromatin status of haploid in hybrid mice is highly correlated with its parent-of-origin (Fig 3A), indicating a genetic or epigenetic inheritance from parents. Interestingly, we found the correlation coefficient between Fb6 and Fcast is higher than that between B6 and Cast (Fig 3A), potentially suggesting that the A|B compartments in hybrid mice are not only inherited from their parents, but also influenced by the micro-environments in the hybrid mouse cells. We further identified the A|B compartments in B6, Cast, Fb6 and Fcast across the genome. We found about 88% of the compartment bins showed exactly the same states among all the 4 samples, essentially consistent with our expectation (Fig S4A). However, 12% of all compartments (870 compartment bins) exhibited different status between B6 and Cast. We retrieved all the genes located in these 870 compartment divergent bins, in which olfactory transduction, neuroactive ligand-receptor interaction, and G-protein coupled receptor signaling pathway were significantly enriched (Fig 3B). The differences of compartment status are potentially associated with significant environmental differences that B6 and Cast have experienced. Indeed, a recent study by Lilue *et al.* found that olfactory clusters are also significantly enriched in the highly different genetic variations among mouse strains (Lilue et al. 2018). Furthermore, it is reported that olfactory-family genes tend to be enriched in the same topologically associated domains (TADs) and be co-regulated (Dixon et al. 2015). We found that TADs, harboring totally more than 33 olfactory clusters, displayed divergent A|B compartments between B6 and Cast. The compartment status in Fb6 and Fcast was the same as that of their parents, respectively (Fig S4B).

**Figure 3.**
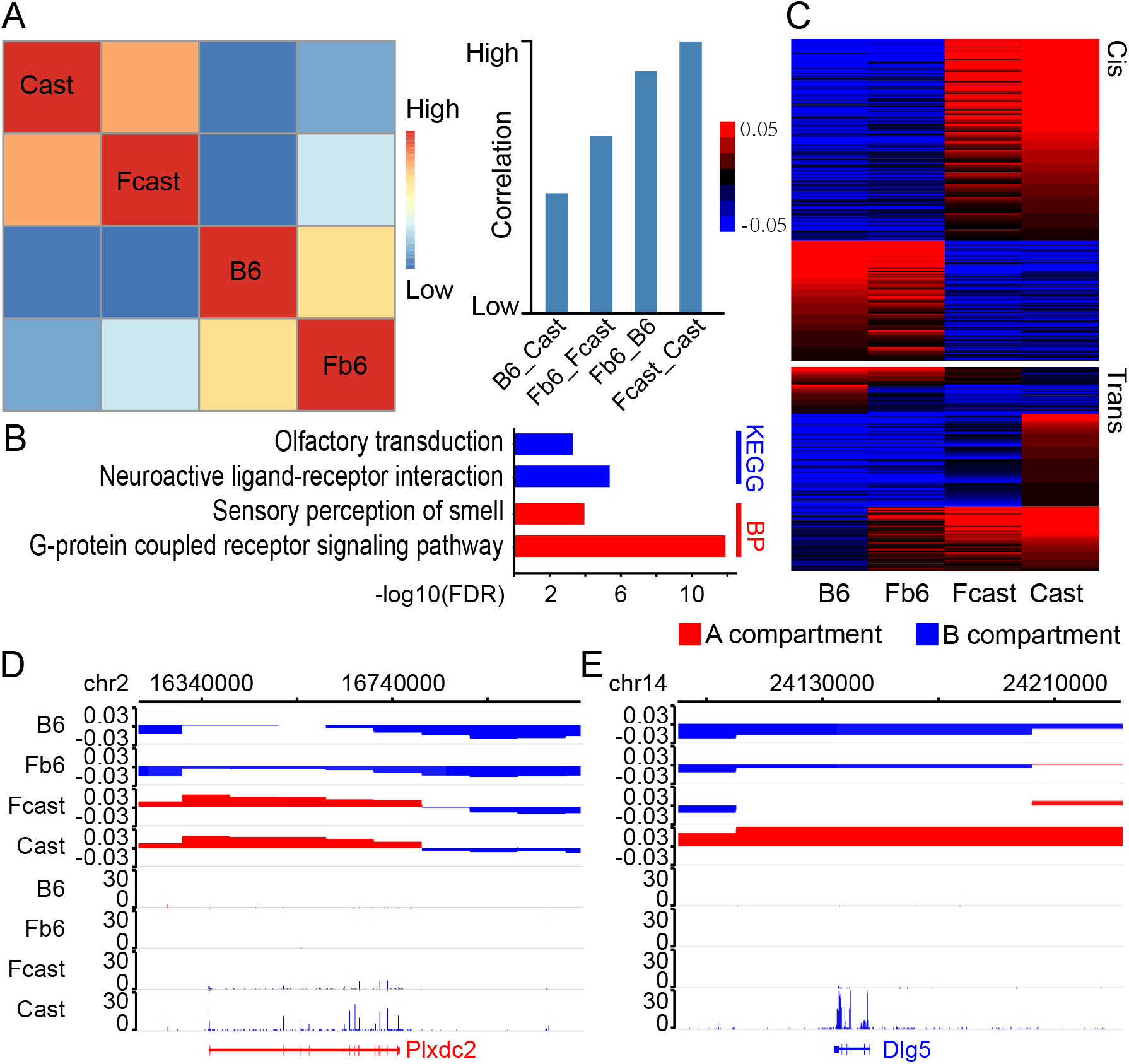
Both cis- and trans-effects of hybrid genomes influenced the status of chromatin compartment and gene expression. A) Correlation of genome-wide A|B compartment scores among the parents and the haploids from hybrid mouse. Heatmap (left) and barplot (right) of correlation coefficient between different genome. B) GO analysis of genes in B6/Cast divergent A|B compartments. C) A|B compartment status of Fb6, Fcast, B6 and Cast in B6/Cast divergent A|B compartment regions. Cis: compartment status is consistent between the haploid genome and parent-of-origin. Trans: compartment status changed in the hybrid mouse. D) A|B compartments and gene expression levels in B6, Cast, Fb6 and Fcast around cis-regulated Plxdc2. E) A|B compartments and gene expression levels in in B6, Cast, Fb6 and Fcast around trans-regulated Dlg5.

For all the 870 A|B compartment divergent bins between B6 and Cast, 58% of the haploid compartments in hybrid mice retained same status as the parent-of-origin. For example, Plxdc2 is located in A|B compartment divergent regions, with A compartment in Cast and B compartment in B6 (Fig 3D, upper tracks). Meanwhile, Plxdc2 is highly expressed in Cast while it is not expressed in B6 (Fig 3D, lower tracks). In this way, the compartment status and gene expression of Plxdc2 in Fb6 and Fcast are consistent with their parents-of-origin (Fig 3D). However, we also found that Fb6 and Fcast display different compartment status compared to their parents-of-origin in 42% of A|B compartment divergent bins. Among the compartments with status changes, 95% converged into same compartment status in hybrid mice (Fig 3C). For example, Dlg5 is associated with inflammatory bowel disease and Crohn disease (Buning et al. 2006; Newman et al. 2006). We found that Dlg5 is located in a compartment divergent region, where it exhibits A compartment and is highly expressed in Cast, while displays B compartment and is not expressed in B6 (Fig 3E). The Dlg5 retains B compartment and is silent in Fb6, while Fcast does not inherit its parental A compartment status in Cast. Instead Dlg5 become B compartment status and is silent in Fcast (Fig 3E).

### TAD boundary shift is associated with gene expression change

TAD is highly self-interacting genomic region, representing the functional and organizational unit of 3D genome (Dekker and Heard 2015; Sexton and Cavalli 2015; Wang et al. 2015; Dixon et al. 2016; Rowley and Corces 2016). Although many methods were developed to identify TADs (Dixon et al. 2012; Filippova et al. 2014; Crane et al. 2015; Durand et al. 2016; Shin et al. 2016; Wolff et al. 2018), it is still inconvenient to quantitatively compare the TADs or TAD boundaries between samples. Here, we introduced the local boundary score (LBS), which is a quantitative value along the genome and facilitates the comparisons between samples (Fig 4A). LBS could be calculated at any scale and any resolution, with peaks being TAD boundaries (see Methods). TAD boundaries and the size of TAD inferred by our method were essentially consistent with that inferred by HiCExplorer (Wolff et al. 2018) (Fig S5A-S5D). We can directly identify the shift of TAD boundaries according to differences of LBS peaks among samples (Fig 4B). Using this approach, we identified a total of 648 TAD boundaries that have exhibited a shift between B6 and Cast (Fig 4C). Genes near the strain-specific boundary showed much higher expression changes than those near the conserved boundaries (Fig 4D).

**Figure 4.**
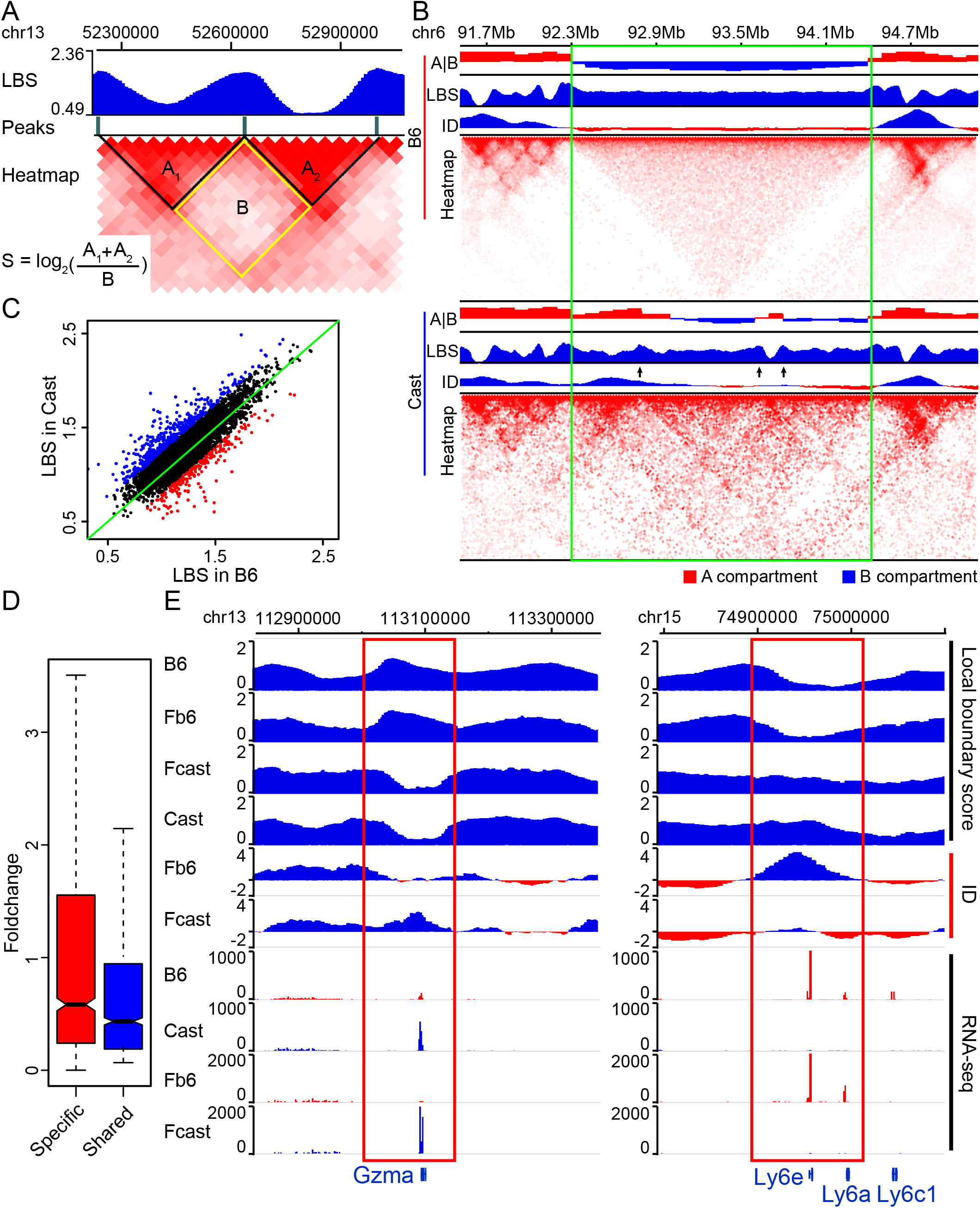
Strain-specific TAD boundary is associated with the expressions of its nearby genes. ID: interaction density. A) Definition and calculation of local boundary scores (LBSs). B) LBSs could accurately detect the TAD boundaries. Strain specific boundaries are associated with chromatin status change and A|B compartment divergence. Green box outlines a region with several strain-specific TAD boundaries (black arrows) and divergent A|B compartments; C) Significantly different TAD boundaries between B6 and Cast inferred by LBS. D) Genes near strain-specific TAD boundaries showed stronger strain-specific expression than these near strain-shared TAD boundaries. Unique: strain-specific TAD boundaries; Conserved: strain-shared TAD boundaries. E) Strain-specific boundaries are associated with strain-specific gene expressions near Gzma and Ly6e, respectively. Red box highlights strain-specific boundary.

Notably, a low LBS means high density of local interactions, hence nearby genes tend to have high gene expression. For instance, two important loci involved in immune function, namely Gzma and Ly-6 family genes (including Ly6e, Ly6a etc.), are located on the shifted TAD boundaries. The Gzma locus in both B6 mice and Fb6 allele is associated with a strong TAD boundary and the Gzma gene is not expressed in both B6 and Fb6 (Fig 4E). In contrast, the Gzma locus is associated with a very low LBS and the gene is highly expressed in both Cast and Fcast (Fig 4E). Similarly, Ly6e locus is not in a TAD boundary and Ly6e is expressed in both B6 and Fb6, while Ly6e locus is in a TAD boundary and ly6e is not expressed in both Cast and Fcast (Fig 4E). Interestingly, LBS within A compartments exhibits substantial fluctuations, while LBS in B compartments is flat and with low variation (Fig S5D). LBS values of the A compartments showed much higher inter-compartment variations than that of B compartments (Fig S5E), indicating both intra- and inter-compartments heterogeneities of A compartments are much higher than that of B compartments. Further analysis showed that one A compartment contained multiple TADs while one B compartment usually contained only one TAD, further indicating the complexity of A compartments (Fig S5F). The multiple TADs within a single A compartment may explain the high variation of LBS values within A compartment.

### Both genetic and epigenetic factors shaped gene expression in hybrid mice

We identified 721 B6 specific and 1,157 Cast specific genes by analyzing RNA-seq data of B6 and Cast naïve CD4 T cells. GO analyses showed that genes associated with immune function are significantly enriched in these strain-specific genes (Fig 5A), consistent with a recent report that environmental adaptation related-genes were significantly enriched among highly diverse genetic regions among mouse strains (Lilue et al. 2018). We further identified 158 Fb6 specific and 178 Fcast specific genes in the hybrid mice, and 72% and 66% of them are overlapped with B6 and Cast specific genes, respectively (Fig S6A). This result indicates that the majority of haploid specific genes show consistent expression with their parental origins, which may be due to cis-effect. The number of differentially expressed genes identified between Fb6 and Fcast is much less compared with that identified between B6 and Cast, largely due to limited allelic reads coverage in Fb6 and Fcast as we only count the reads containing strain specific polymorphisms.

**Figure 5.**
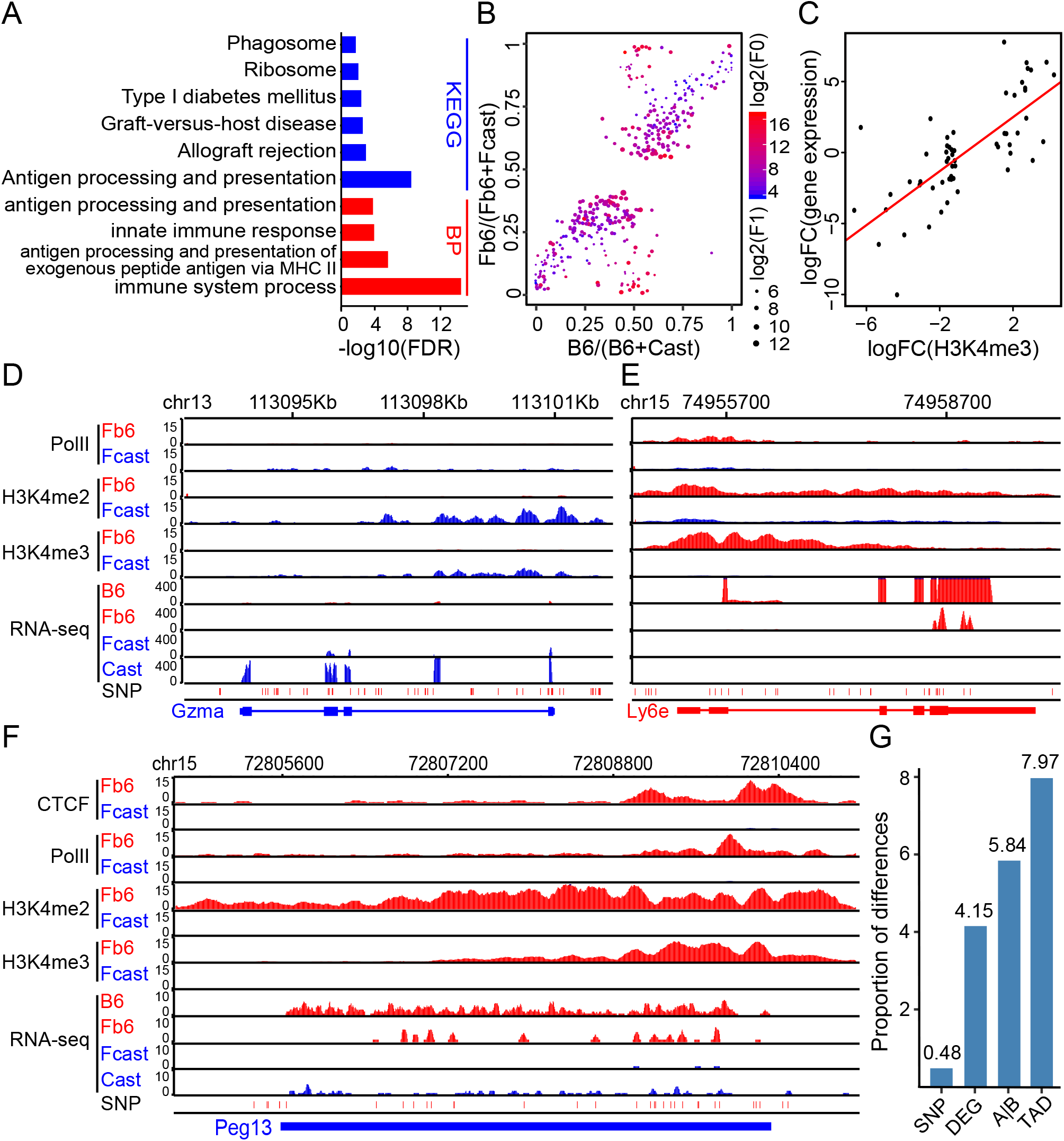
Both genetic and epigenetic regulations shaped the gene expression in hybrid mouse. A) Gene ontology enrichment for differentially expressed genes between B6 and Cast. B) Relative gene expressions in parental mice and two haploids of differentially expressed genes between Fb6 and Fcast. Dot size represents average expression level of Fb6+Fcast (F1). Blue to red represents average expression level of B6+Cast (F0). C) Allele-specific H3K4me3 is positively correlated with allele-specific gene expression in hybrid mouse. Each point represents a biased ChIP-seq peak and its regulated gene. (R = 0.85, P value = 1.2e-13) D) Cast-and Fcast-specific epigenetic modifications near Gzma are associated with Gzma specific expression. E) B6-and Fb6-specific epigenetic modification near Ly6e are associated with Ly6e specific Ly6e expression. F) Maternal imprinted gene Peg13 only exhibited paternal specific active epigenetic modifications in hybrid mouse. G) Fraction of differences between B6 and Cast at genomic, transcriptomic and epigenomic level.

In order to gain insights into the regulation of differentially expressed genes between Fb6 and Fcast, we displayed their relative gene expression levels in parental mice and two haploids in one plot. The genes distributed along the diagonal were consistently differentially expressed in both parental mice and two haploids of the hybrid mice (Fig 5B), which can be mainly attributed to cis-effects. The genes distributed near the vertical line were differentially expressed in the two haploids of the hybrid mice while not differentially expressed in the parental mice (Fig 5B), which may be attributed to trans-effects. Thus, both cis- and trans-effects played an important role in gene regulation in the hybrid mice, consistent with a study using liver tissues (Goncalves et al. 2012). After separating the ChIP-seq reads from hybrid mice into Fb6 and Fcast, similar to the analysis of Hi-C data, we found the allele-specific H3K4me3 and H3K4me2 modifications are positively correlated with allele-specific gene expression (Fig 5C, R = 0.85, P value = 1.2e-13; and S6B, R = 0.82, P value < 2.2e-16). As expected, Gzma and Ly6e showed accordant allele-specific histone modifications with the allele-specific gene expression in the hybrid mice (Fig 5D-5E), suggesting that cis-effects could be related to either genetic or epigenetic mechanisms. Paternally Expressed 13 (Peg13), associated with Birk-Barel mental retardation dysmorphism syndrome (Barel et al. 2008; Court et al. 2014), is a trans-effect gene identified in the hybrid mice and has been reported as maternal imprinted gene (Smith et al. 2003). Although Peg13 expression was detected in both Cast and B6, it is silent in Fcast (maternal) and it has Fb6 (paternal)-specific active histone modifications, CTCF binding and expression (Fig 5F), potentially indicating imprinted gene showed trans-effect pattern in hybrid mice.

### Epigenetic differences significantly contribute to B6 and Cast difference

Although it is well known that genetic change is fundamental force of evolution (Gandon and Nuismer 2009; North et al. 2011), epigenetic variations also play an important role in evolution and phenotype change (Romero et al. 2012; Schmitz et al. 2013; Gutierrez-Arcelus et al. 2015). Thus, both genetic variations and epigenetic variations could explain the phenotypic difference between B6 and Cast. We calculated the difference between B6 and Cast in each type of variation, which may contribute to the adaption of mouse strains to their local environments. We found epigenetic differences between B6 and Cast are much higher than that of genetic difference (Fig 5G). Genetic difference between B6 and Cast only account less than 1% genomic sequence, while more than 5% A|B compartments status and TAD boundaries are different between B6 and Cast. The levels of epigenetic differences are comparable to the level of differentially expressed genes between B6 and Cast (5%), indicating that epigenetic factors may be the major contributors of gene expression difference between B6 and Cast. The epigenetic difference between B6 and Cast could explain the large fraction of differentially expressed genes even though there is only a small fraction of genetic difference between B6 and Cast.

## Discussion

How the 3 billion base-pair genomic DNA is organized in the nucleus has been a subject of extensive research. Previous imaging studies based on chromosome paintings elegantly demonstrated that each chromosome exists in a space termed chromosome territory (Cremer et al. 2003; Bolzer et al. 2005; Hua and Mikawa 2018). However, it appears that the positioning of chromosome territories, particularly the relative position of homologous chromosomes, vary dramatically across different cells. Due to the relatively small number of cells that can be examined by this kind of techniques, it has been difficult to reach a consensus about the relative position between homologous chromosomes within a nucleus. In this regard, the Hi-C methods provide information on the 3D nuclear structure by averaging millions of cells (Lieberman-Aiden et al. 2009; de Wit and de Laat 2012; Denker and de Laat 2016). However, previous studies lacked sufficient allele-specific information to resolve the two homologous chromosomes and thus only inferred average interaction profiles of homologous chromosomes. A recent single-cell Dip-C study explored the naturally existing SNPs in human cells and observed interesting differences between the maternal and paternal alleles (Tan et al. 2018). In this study, we utilized the 20 millions of SNPs existing between B6 and Cast (Keane et al. 2011; Yalcin et al. 2012), enabling assigning the Hi-C, ChIP-seq, and RNA-seq sequence reads to strain specific haploids at high-resolution. To some extent, this study was the first endeavor that focuses on investigation of 3D nucleus organization of each haploid chromosome.

Previously, B6 and Cast hybrid mice had been used for investigation of cis- and trans-effect of gene regulation and the B6 haploid and Cast haploid could be distinguished from each other based on strain specific alleles (Goncalves et al. 2012). However, the allele specific epigenetic regulation is rarely studied and dynamics of epigenetic status in hybrid mice is unknown. Interestingly, haploids with divergent compartments converged into the same A|B compartment in hybrid mice due to shared cellular environment. The gene expression pattern can be changed in the hybrid cells due to cis- and trans-effects (Fig 5B), along with the transition of histone modifications and binding of transcription factor on imprinted genes (Fig 5F), further confirming that micro-environment plays a critical role in the epigenetic regulation on gene expression.

Our 3e Hi-C data indicated that the inter-chromosomal interaction pattern is highly similar between homologous chromosomes. Furthermore, we found that the positioning of homologous chromosomes in 3D nucleus is symmetric to the center of the nucleus. Since homologous chromosomes have similar interaction patterns and form similar local environments, the best model to fit two differently located objects with similar local environments should be a symmetric model. Most importantly, symmetric positioning of homologous chromosomes has many implications for understanding gene regulation and nucleus organization. For example, although many studies inferred the TADs, A|B compartments and chromatin loops using averaged interaction matrix from different cells and from different homologous chromosomes, the conclusions from these studies are reasonable to some extent; even though these studies did not consider the difference between the two haploids because the homologous chromosomes showed much similar interaction patterns.

In summary, our study indicates symmetrical positioning of homologous chromosomes in the 3D nuclear space and provides allele-specific histone modification and RNA PolII binding profiles that are important for understanding the epigenetic regulation of differential gene expression between homologous chromosomes.

## Methods

### Mice and Isolation of Cells

Both C57BL/6J (B6) mice and CAST/EiJ (Cast) mice were purchased from Jackson Laboratories (Bar Harbor, ME, USA). This study was reviewed and approved by the Animal Care and Use Committee of the National Heart Lung and Blood Institute. All mice received humane treatment according to NIH guidelines and the ‘‘Guiding Principles for Research Involving Animals and Human Beings.’’

Male B6 and female Cast mice were crossbred to generate the F1 hybrid mice. Naïve CD4 T cells were purified from spleen and lymph nodes of B6, Cast and hybrid mice with CD4 microbeads (Miltenyi Biotech, CD4^+^CD62L^+^ T cell isolation kit II, mouse). The purity of naïve CD4 T cells was assessed by flow cytometry for CD4^+^, CD8^−^ and CD62L^+^ using FACSCanto II (BD Biosciences). Over 98% purity of CD4^+^CD8^−^CD62L^+^ cells was considered for further experiments. All experiments were conducted on 6 to 8-week-old mice. Antibodies including anti-CD4 (RM4–5) and anti-CD62L (MEL-14) for flow-cytometry analyses were purchased from eBiosciences. Dead cells were excluded by DAPI staining.

### 3e Hi-C

The multiple-enzyme Hi-C (3e Hi-C) was performed according to our previous studies (Ren et al. 2017; Hu et al. 2018). About 1,000,000 naive CD4 T cells were cross-linked with 1% formaldehyde for 10 mins. Cells were lysed and digested with 20 Units CviQ I (NEB), and 20 Units CviA II (NEB) at 25°C for 20 minutes, then 20 Units Bfa I (NEB) at 37°C for 20 minutes. The reaction was stopped by washing the cells twice with 600 mL wash buffer (10mM NaCl, 1mM EDTA, 0.1% triton-100). DNA ends were marked by biotin-14-dATP with Klenow (large) for 1h at 37°C. Blunt-end DNA fragments were ligated with T4 DNA Ligase overnight at 16°C. DNA was then reverse cross-linked and purified by phenol/chloroform extraction. Biotin was removed from unligated DNA-ends by T4 DNA polymerase for 2hs at 12°C. DNA was purified by phenol/chloroform and sheared to 300-500 bp by sonication followed by DNA-end repair and addition of adenosine. Biotin labeled DNA was pull-downed by streptavidin beads followed by Illumina adaptor ligation and PCR amplification. DNA fragments of 300 to 700 bp were isolated 2% agarose gel and sequenced by paired-end sequencing on Illumina Hiseq 2500.

### ChIP-Seq

ChIP-Seq experiments were performed as described previously (Barski et al. 2007). In brief, 1,000,000 of Naïve CD4 T cells were fixed for 10 min with 1% formaldehyde in complete medium, sonicated and chromatin immunoprecipitation was performed with antibodies against CTCF (07-729, Millipore), RNA Polymerase II (ab5408, Abcam), H3K4me3 (ab8580, Abcam) and H3K4me2 (ab32356, Abcam). The DNA was then ligated with the ‘Y’ shaped Illumina adaptor and amplified for 18 cycles using indexing primers as described. PCR products between 160-300bp were isolated on 2% E-gel for sequencing on Illumina Hiseq2500.

### RNA-Seq

Total RNA was extracted and purified with miRNeasy micro kit (217084; Qiagen) and DNase set (79254; Qiagen), followed by delusion with 10 mL of RNase-free water. Purified total RNA was reverse transcribed with the Ovation RNA-Seq System V2 (7102-08; NuGEN Technologies). cDNA was sonicated in a Diagenode Bioruptor (level M, for a total of 30 min of 20s on and 20s off) to obtain fragment in the size range of 100-400bp. Indexed libraries were prepared with a Multiplexing Sample Preparation Oligonucleotide Kit (1005709; Illumina) and DNA End-repair Kit (ER81050; Epcentre) according to the user’s manual (Epcentre) and sample preparation guide (Illumina).

### SNPs calling and cross validation

We downloaded the files containing mouse indels and SNPs information (about 21 million SNPs) (mgp.v5.merged.indels.dbSNP142.normed.vcf) from Mouse Genomes Project (ftp://ftp-mouse.sanger.ac.uk) (Keane et al. 2011; Yalcin et al. 2012) and examined the SNPs between C57BL/6J and CAST/EiJ using SNPsplit toolkit (Krueger and Andrews 2016). Next, we built the mm10 Bowtie2 index with all SNP positions being masked by the ambiguity base ‘N’. All reads from B6 and Cast Hi-C libraries were mapped to this N-masked reference using Bowtie2 (Langmead and Salzberg 2012) (with parameter --very-sensitive -L 30 --score-min L,-0.6,-0.2 --end-to-end). For retrieving the highly confident SNPs, we re-called the genotype for each SNP locus in B6 and Cast Hi-C libraries using mpileup in samtools (Li et al. 2009) with reads MAPQ >= 30. The called SNPs showing identical states with that downloaded from mouse genome project were kept for further analyses. In total, we obtained 13 million highly confident SNPs for distinguishing Cast haploid reads and B6 haploid reads in hybrid mice.

### Hi-C, ChIP-seq and RNA-seq data processing

The paired-end reads from Hi-C, also known as paired-end tags (PETs), were mapped to N-masked mm10 reference with bowtie2 separately. Both ends of the PETs with MAPQ >= 30 were kept for further analyses. The PETs with the same start and the same end were treated as redundant PETs, among which only one PET was kept for further analysis. Intra-chromosomal PETs within 10Kb were considered as self-ligations and were filtered out. The interaction matrices were constructed at different scales, with bin sizes varying from 20Kb to 100Kb. The matrices were normalized using iterative correction algorithm ICE (Imakaev et al. 2012), and further optimized in HiC-Pro (Servant et al. 2015). A|B compartments were calculated as described in Lieberman-Aiden et al (Lieberman-Aiden et al. 2009). PETs from hybrid mice Hi-C library were split into haploid using SNPsplit using highly confident SNPs, and the downstream processing was similar to normal Hi-C samples.

ChIP-seq reads were mapped and filtered similarly to Hi-C reads, and extraction of reads for generation of haploid ChIP-seq data was similar to the generation of haploid Hi-C. Peaks were called using macs2 (Zhang et al. 2008). For comparison of the differences of histone modifications and TF bindings between Fb6 and Fcast, we counted the allele-specific reads located in the peaks using *intersect* in bedtools (Quinlan 2014). The significances of the differentially binding sites were calculated by DESeq (Love et al. 2014). ChIP-seq reads were extended to 150bp and normalized by sample size, then converted to bedGraph for visualization.

RNA-seq reads were mapped to N-masked mm10 reference by TopHat2 (Kim et al. 2013), and reads with MAPQ >= 30 were kept for further analyses. The batch effect of replicates was adjusted by ComBat from R packa ge sva (Leek et al. 2012), and differentially expressed genes were identified by DESeq2 (Love et al. 2014) with FDR < 0.05 and fold change > 1.8. Allele-specific reads in turn were grouped by SNPsplit, whereas allele-specific gene expression was counted and quantified as normal.

### PCA analysis of inter-chromosomal interaction matrix

In order to analyze the inter-chromosomal interaction patterns in the nucleus of hybrid mouse, we retrieved all the inter-chromosomal PETs with both ends containing confident SNPs in hybrid mouse Hi-C data. We generated chromosomal resolution interaction matrix for all the 40 paternal and maternal chromosomes. To eliminate the bias induced by chromosome size, the PETs number was normalized by formula (1) or ICE normalization (Imakaev et al. 2012). We found the first two PCs contribute majority of inter chromosomal variations after we conduced PCA on the allelic specific interaction matrix. In order to illustrate the different interaction patterns between active region and inactive region on X chromosome, the full-length X chromosome was separated into X-a and X-i at the Dxz4 loci. To remove the effects from interactions between homologous chromosomes, the interaction counts between homologous chromosomes were set to zero when conduct PCA.

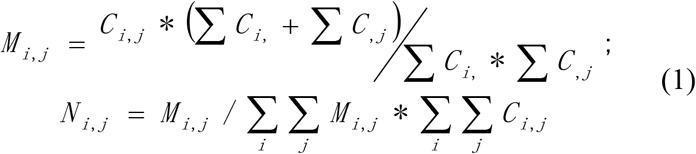

Where *C_i,j_* is the raw interaction count for each chromosome pair i and j, *M_i,j_* is the intermediate count after normalization and *N_i,j_* is the final normalized count which is scaled to the original sample size.

### Modeling the 3D nucleus in hybrid mouse

For construction of 3D nucleus of hybrid mouse, we developed an iteratively weighted adjusting algorithm to infer the relative positioning of each chromosome in 3D space. The purpose of iterative process of this model is to continuously minimize the sum of errors between coordinate-based distance and the ‘real’ distance which was converted from the allele-specific interaction matrix (Fig S1A). In brief, we first initialized random xyz values for each chromosome, then iteratively adjusted the xyz values based on the distance errors between that chromosome and the other chromosomes. We noted that this model is more likely to reach the local optimum and steady state after finite iterations (Fig S3A). The mathematical formulae of this model is:

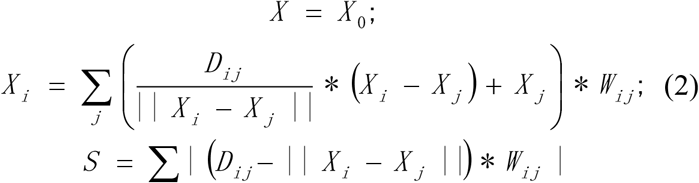

Where *X_0_* is randomly initialized 3D coordinates (xyz values) for each chromosome, *D_ij_* is the ‘real’ distance matrix converted from the allele-specific interaction matrix based on the fitted PETs count to distance function (Fig S1A), *W_ij_* represents the weight matrix for each chromosome pair converted from the allele-specific interaction matrix. The aim of the iteration is to achieve the smallest *S*.

Since the interaction between homologous chromosomes may introduce bias to the 3D model, we reset the interactions between homologous chromosome pairs to the average interaction density of the corresponding chromosome to all non-homologous chromosomes. We also constructed the 3D model with split active and inactive X chromosome (Fig S3C) and without X chromosomes (Fig S3D).

### Local boundary score (LBS) analysis

LBS is defined as the logarithm ratio of intra-loci interactions to inter-loci interactions in a given local region (Fig 4A). In details, genome was separated into many small bins with equal size and several continuous bins made up a locus (for example, bin size=2k and locus=150k). For a given bin, the interactions within its left locus (A_1_ in Fig 4A) and its right locus (A_2_ in Fig 4A) are defined as its intra-loci interactions. The interactions between these two loci (150kb x 150kb, B in Fig 4A) are inter-loci iterations, then the log2-transformed ratio of intra-loci interactions to inter-loci interactions was calculated as LBS of the bin. To avoid false positive in sparse interactions, chromosome-wide average value was set as the initial background for intra- and inter-loci interactions. In this way, the peak of LBS is the relative inter-loci interaction in local region reaches the maximum value, which indicated the presence of the TAD boundary. Peakdet (http://billauer.co.il/peakdet.html) was used to call local peaks of LBSs. To compare TAD boundary shift between samples, LBS biases were calculated using ROSE that initially for distinguishing super enhancers from typical enhancers (Whyte et al. 2013).

We compared the TADs called by LBSs with that by HiCExplorer (Wolff et al. 2018), with default parameters. Results showed that more than 80% of the TAD boundaries called from two methods were exactly the same (Fig S5B), and the median sizes of TADs at the given resolution are 300Kb for both methods (Fig S5C), indicating that LBS is efficiently for the identification of TADs at very high resolution.

## Data access

All sequencing data have been deposited in Gene Expression Omnibus (GEO) under accession number GSE132898.

## Acknowledgements

We thank the National Heart, Lung, and Blood Institute DNA Sequencing Core Facility for sequencing the libraries; and the National Heart, Lung, and Blood Institute Flow Cytometry Core facility for sorting the cells. The work was supported by Division of Intramural Research, National Heart, Lung and Blood Institute (K.Z.), National Key R&D Program of China (2018YFC1004500) (W.J.), National Science Foundation of China (81872330, 31741077) (W.J.) and Shenzhen Innovation and Technology Commission (JCYJ20170817111841427) (W.J.).

## Author Contributions

K.Z. and W.J. conceived this project. K.C. and K.P. did the experiments. Z.H. performed the data analysis. W.J. and K.Z. supervised this project, with contribution from W.C., N.H. and C.L.. Z.H., W.J. and K.Z. wrote the manuscript with inputs from all authors.

---

**Figure S1.**
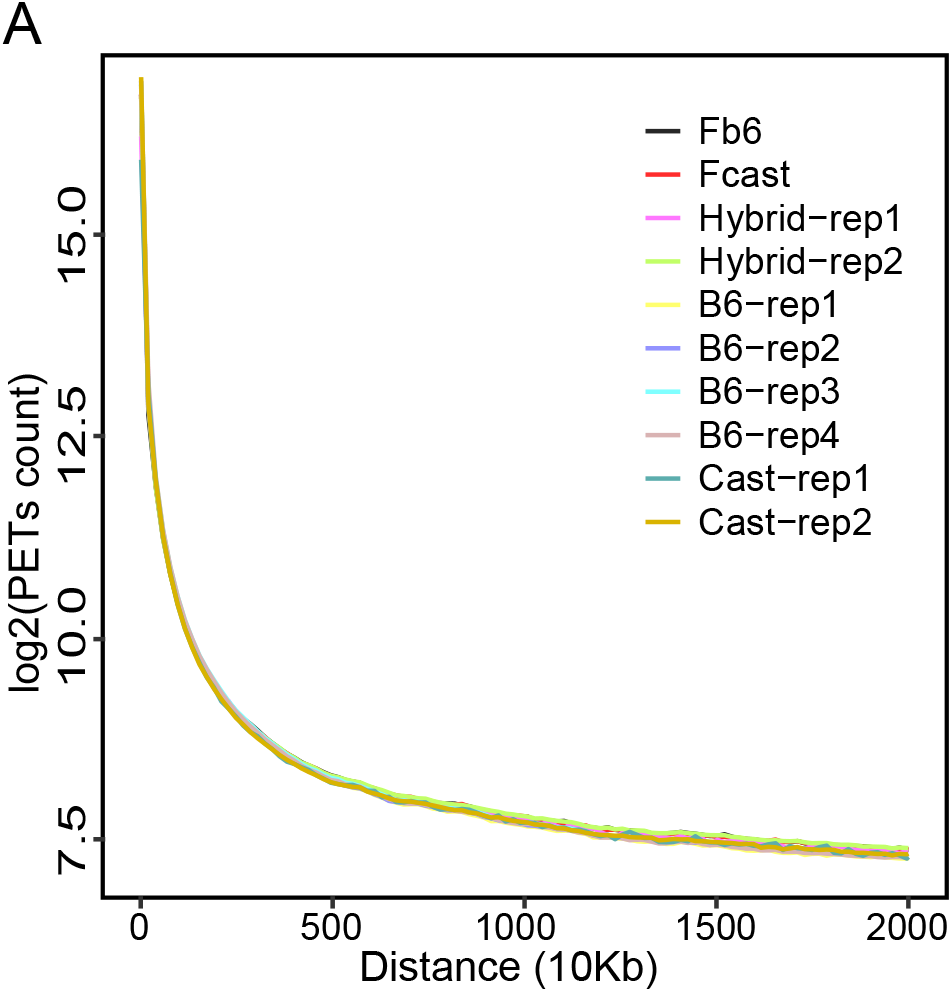
PETs density along distance for Hi-C replicates and haploid in hybrid mouse.

**Figure S2.**
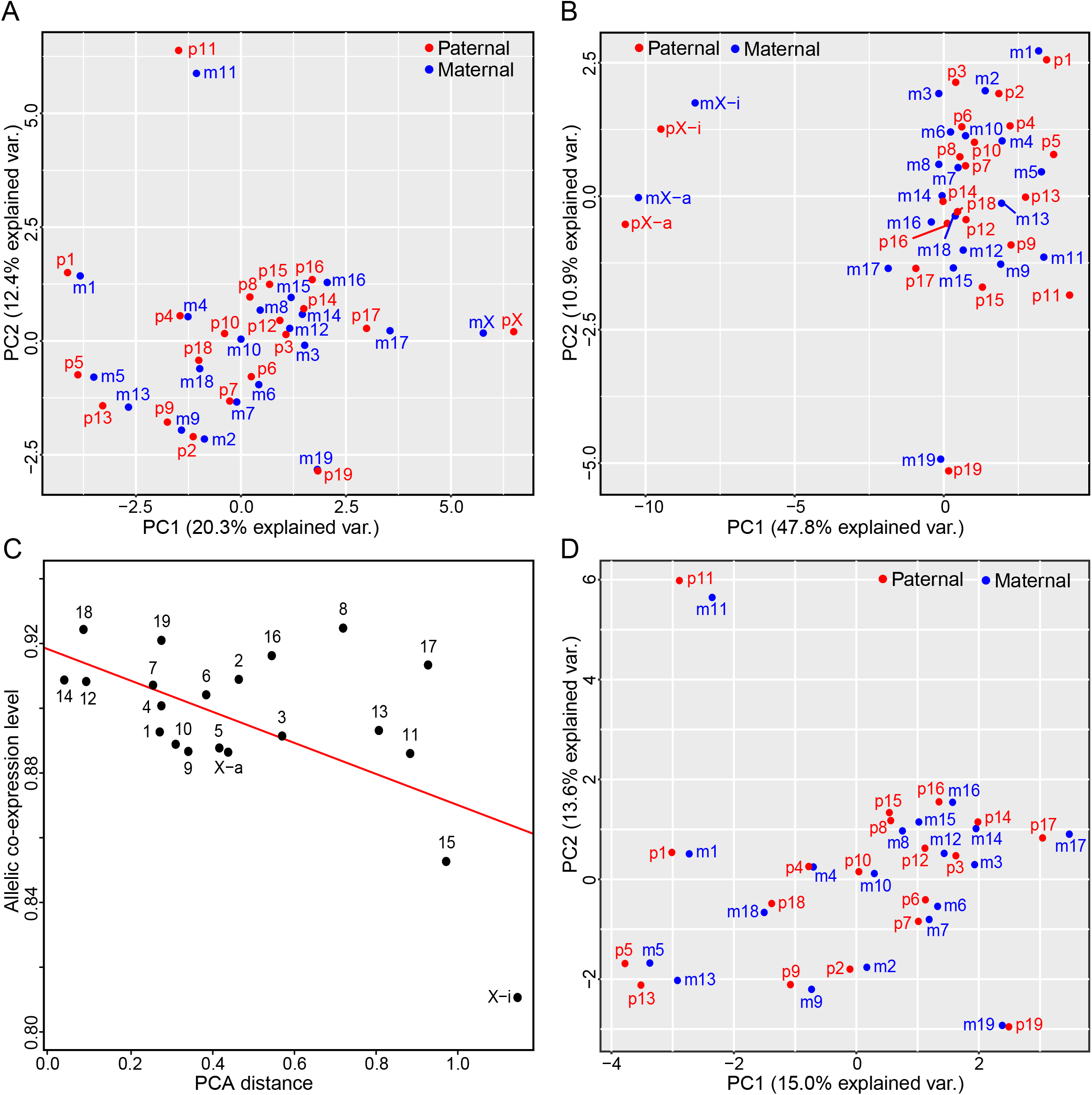
Homologous chromosomes show similar interaction patterns. A) PCA analysis of chromosomal interactions by excluding PETs between homologous chromosomes. B) PCA analysis of chromosomal interactions with X chromosome being separated into active and inactive regions. a: active region; i: inactive region. C) Allelic co-expression between homologous chromosomes is correlated with the similarity of chromosomal interaction pattern, with X chromosome being separated into active and inactive regions. (R = 0.66, P value = 0.001) D) PCA analysis of chromosomal interactions by excluding X chromosomes.

**Figure S3.**
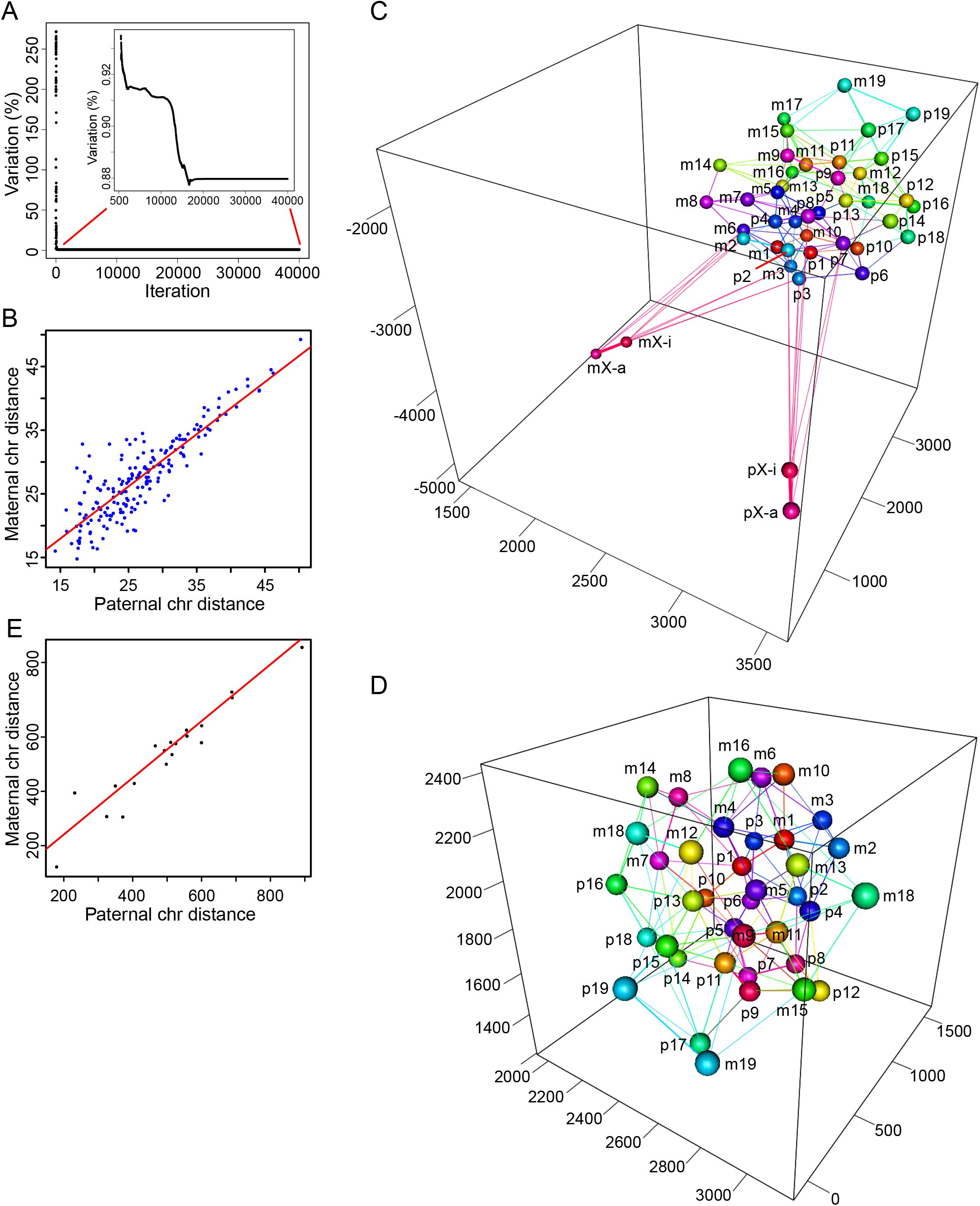
Constructing the 3D nucleus using iteratively adjusting model in the hybrid mouse. A) Variation of the model decreased rapidly during iteration. B) The distances of homologous chromosome pairs to any other chromosome are highly correlated in the 3D model. (R = 0.87, P value < 2.2e-16) C) 3D nucleus when X chromosomes separated into active and inactive regions. a: active region; i: inactive region. D) 3D nucleus without X chromosomes. E) Distance to 3D nucleus center for paternal chromosomes is highly correlated with that of maternal homologous compartment in 3D nucleus without X chromosomes. (R = 0.95, P value = 1.8e-10)

**Figure S4.**
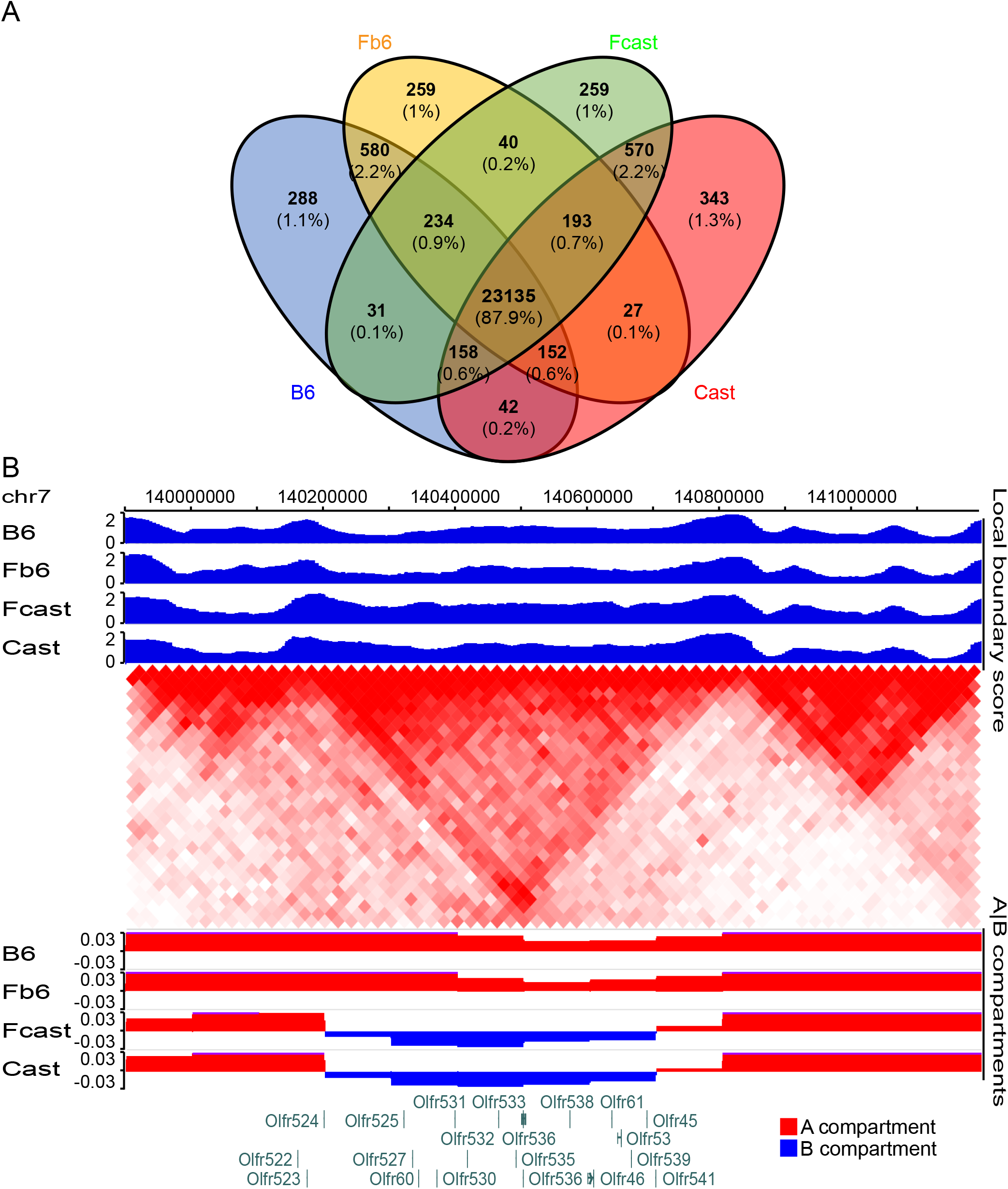
A|B compartment status among B6, Cast and two haploids in hybrid mouse. A) Venn diagram showing significant overlaps of A|B compartment among parents and two haploids of hybrid mouse. B) The olfactory genes are located within a single TAD, with divergent A|B compartment between B6/Fb6 and Cast/Fcast.

**Figure S5.**
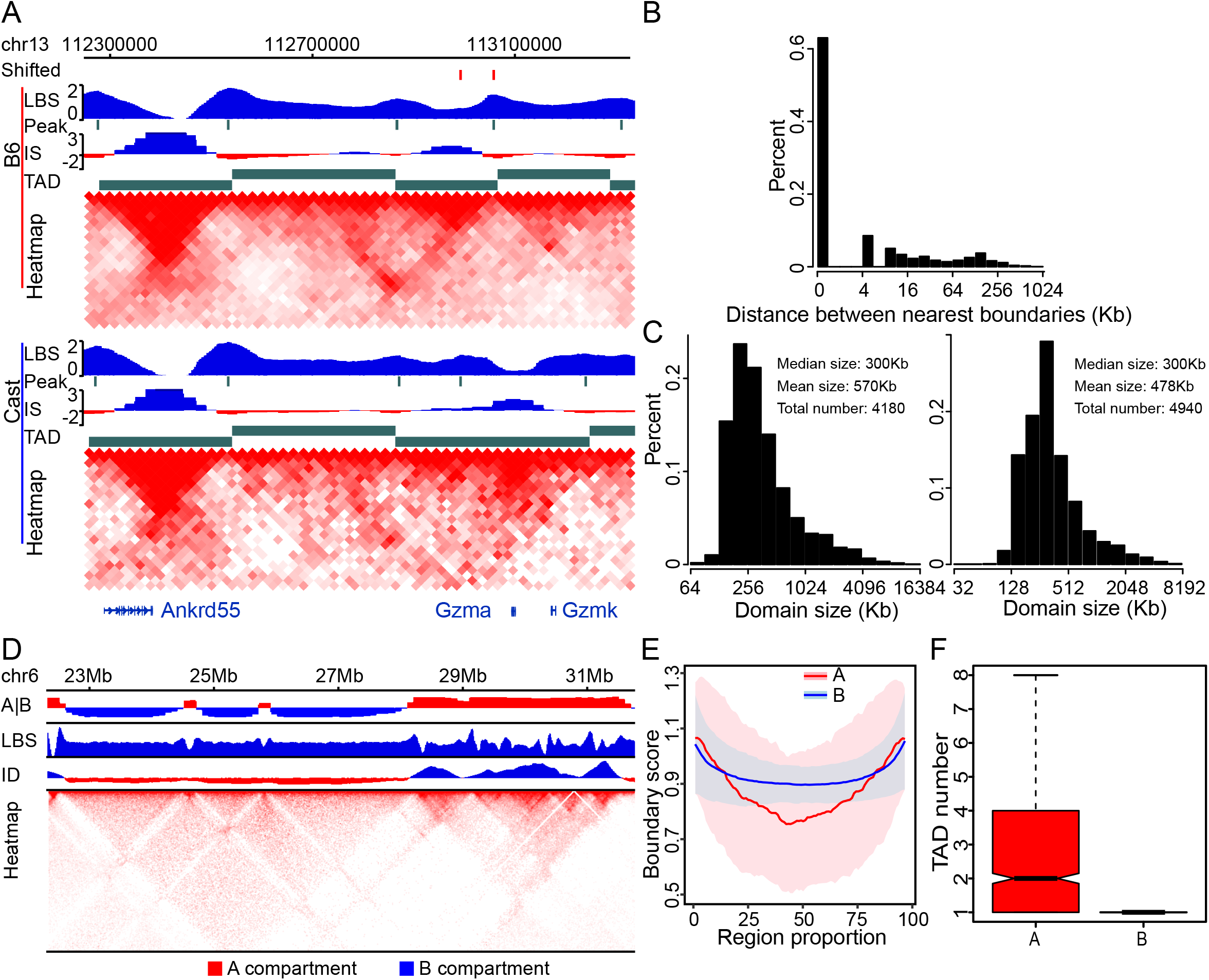
TADs calling by LBS and LBS/TADs differences between A and B compartment. A) TAD boundaries called by LBS and HiCExplorer are highly consistent. Shift: the different TAD boundary between B6 and Cast. Peak: LBS peaks which represent the TAD boundaries. IS: insulation score calculated by HiCExplorer. TAD: TAD called by HiCExplorer. B) Histogram of distance between TAD boundaries called by LBS and these nearest called by HiCExplorer. C) Distribution of TADs size called by LBS (left) and HiCExplorer (right). D) LBS in B compartments are flat and smooth, while LBS in A compartment fluctuate. A|B: A|B compartment; ID: interaction density. E) Comparison of mean and standard derivation (SD) of LBS in A and B compartments, showing a high variability of LBS at A compartment. F) Boxplot of number of TADs in each A or B compartment. (P value < 2.2e-16)

**Figure S6.**
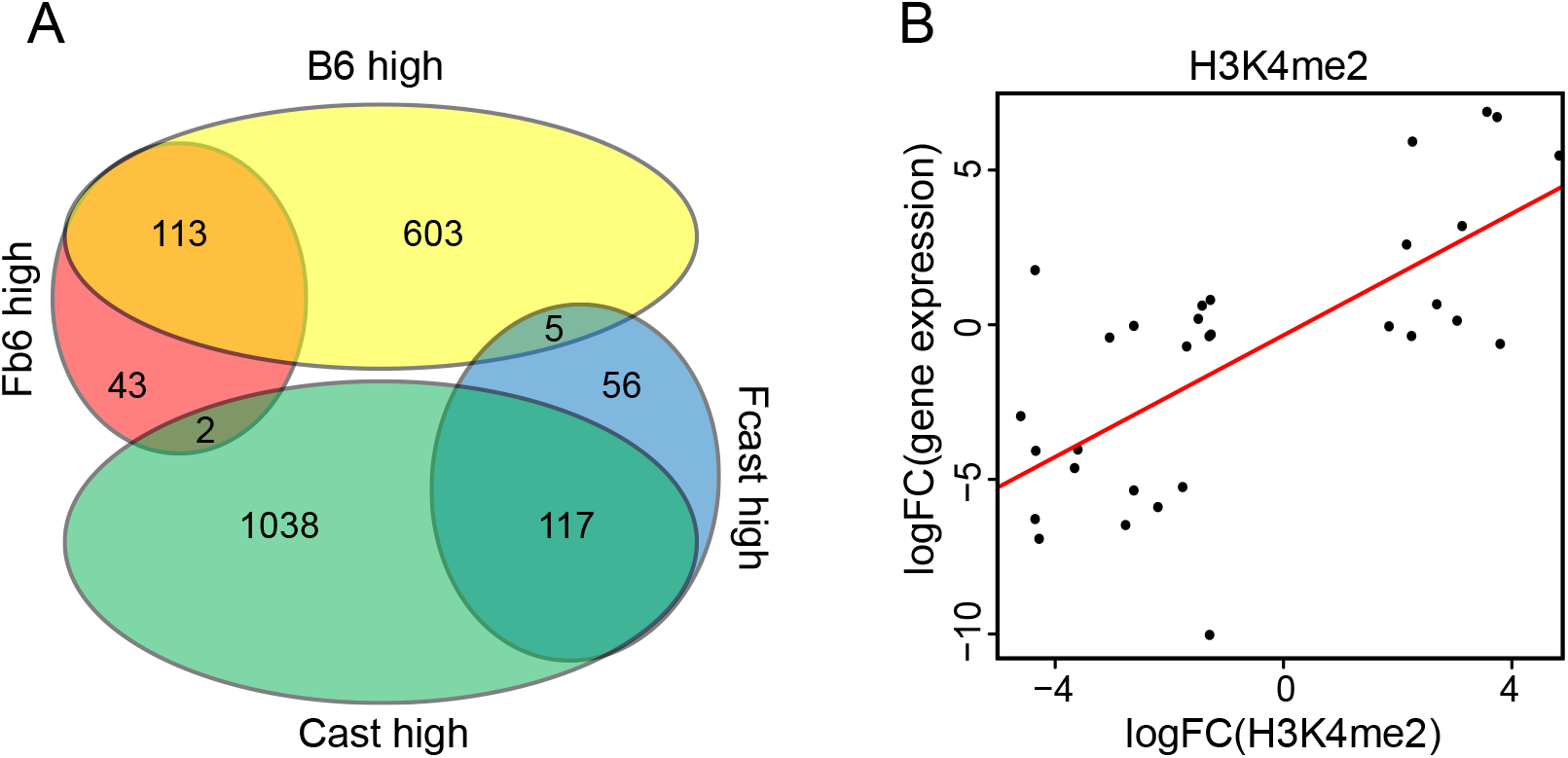
Allelic specific expressed genes and its association with allelic epigenetic modification. A) Venn diagram showing the overlap of DEGs between B6/Cast and Fb6/Fcast. B) Allele specific H3K4me2 is positively correlated with allele-specific gene expression in hybrid mouse. Each point represents a biased ChIP-seq peak and its regulated gene. (R = 0.82, P value < 2.2e-16)

